# Structure-based discovery of Saponarin as a broad-spectrum allosteric inhibitor of banana viral coat proteins

**DOI:** 10.1101/2025.06.30.662402

**Authors:** Mohammad Imran Kumasagi, N. Nagesha, Dyumn Dwivedi, G. M. Santhosh, Shridhar Hiremath, Uddalak Das, K.S. Shankarappa

## Abstract

Banana (*Musa spp*.), a globally significant staple crop, suffers substantial yield losses from persistent viral infections caused by Banana Bunchy Top Virus (BBTV), Banana Streak Virus (BSV), Banana Bract Mosaic Virus (BBrMV), and Banana Mosaic Virus (BMoV). Given the critical role of viral coat proteins (CPs) in genome encapsidation, movement, and host infectivity, these capsid components represent attractive targets for antiviral intervention. Here, we report a comprehensive *in silico* pipeline integrating homology modeling, structure-based molecular docking, pharmacokinetic profiling, and 100-ns all-atom molecular dynamics (MD) simulations to identify potential CP inhibitors from a curated phytochemical library. High-confidence structural models of the CPs were generated using SWISS-MODEL and AlphaFold3 and validated via Ramachandran analysis, ERRAT, and Verify3D. Virtual screening of 100 plant-derived compounds revealed Saponarin, a flavonoid glucoside, as the top-scoring molecule across all viral targets, with docking scores ranging from –13.33 to –8.75 kcal/mol. Binding interactions were dominated by extensive hydrogen bonds and π-based stacking with conserved aromatic and polar residues within the capsid interface pockets. ADMET predictions indicated Saponarin possesses favorable physicochemical properties, high aqueous compatibility, low clearance, and minimal ecotoxicological risk. MD simulations confirmed stable binding, persistent hydrogen bonding, and conserved protein compactness, supporting an allosteric inhibition mechanism. These findings establish Saponarin as a structurally and pharmacologically viable broad-spectrum antiviral candidate for banana virus control, warranting experimental validation for translational deployment in sustainable crop protection strategies.

Graphical Abstract:
Schematic representation of the structure-based virtual screening workflow for identifying broad-spectrum inhibitors targeting coat proteins (CPs) of BBTV, BSV, BBrMV, and BMoV. Homology models of CPs were constructed, followed by binding site prediction, ligand retrieval, and molecular docking of 100 phytochemicals. Saponarin emerged as the top candidate, showing strong binding at conserved allosteric sites. ADMET profiling confirmed favorable pharmacokinetics and low toxicity. Molecular dynamics simulations validated the stability of Saponarin–CP complexes, supporting its potential as a broad-spectrum antiviral agent against multiple banana viruses.

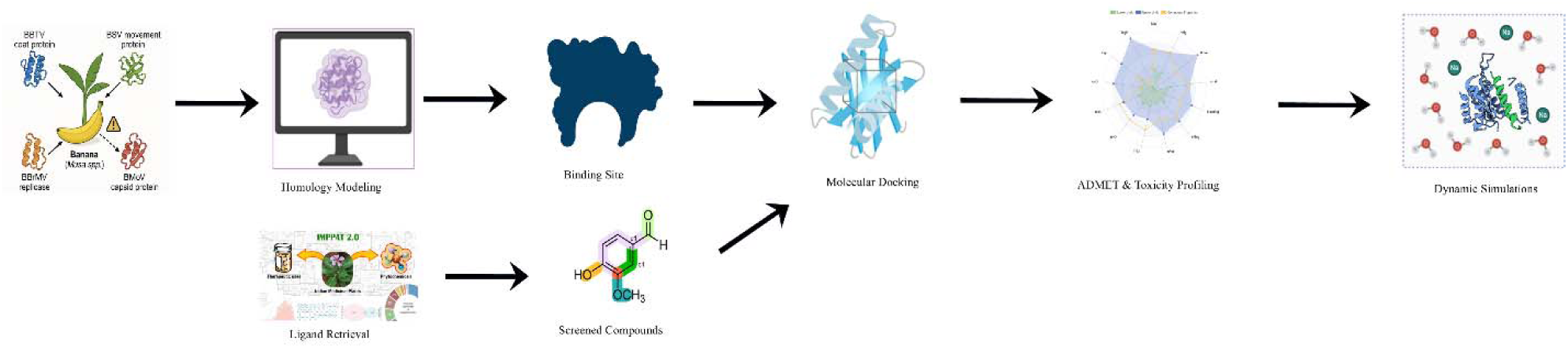

## 1. Introduction

Banana (*Musa spp.*), a staple fruit crop native to Southeast Asia, holds immense global significance both economically and nutritionally. As one of the most consumed fruits worldwide, bananas serve as a primary source of nutrition for over a billion people, particularly in tropical and subtropical regions [1]. Cultivated varieties primarily derive from two wild species, *Musa acuminata* (A genome) and *Musa balbisiana* (B genome), forming diverse genomic groups such as AAB and ABB triploids (2n = 3x = 33) [2]. In India, banana cultivation spans approximately 960 thousand hectares with an annual production of about 33.83 million metric tonnes, predominantly in Tamil Nadu, Andhra Pradesh, Maharashtra, Karnataka, and Gujarat [3].

Despite the crop’s vital role in food security and rural livelihoods, banana production faces severe constraints, primarily due to the emergence and spread of viral diseases. These viral infections are of particular concern owing to their systemic nature, ease of spread through vegetatively propagated suckers, and efficient transmission by insect vectors [4]. Unlike many fungal or bacterial infections, the symptoms of viral diseases in bananas are often subtle or variable, making field diagnosis challenging. The intensification of banana cultivation in recent decades has further exacerbated the incidence and severity of viral outbreaks, leading to substantial yield and quality losses [5].

Among the numerous viruses affecting banana crops, four are of national and international concern: Banana bunchy top virus (BBTV), Banana streak virus (BSV), Banana bract mosaic virus (BBrMV), and Cucumber mosaic virus (CMV). These pathogens are widespread across banana-growing regions and collectively cause significant economic damage.

BBTV, a member of the *Nanoviridae* family and *Babuvirus* genus, is considered the most destructive banana virus. It comprises at least six circular, single-stranded DNA components, each around 1.1 kb, encapsulated within icosahedral virions (∼20 nm). The virus is persistently transmitted by the banana aphid *Pentalonia nigronervosa*. Symptoms include the characteristic “Morse code streaking” on the midrib and petioles, lamina narrowing, and a bushy plant appearance, often resulting in complete yield loss [6,7].

BSV, a *Badnavirus* with a 7.2 kb circular double-stranded DNA genome, spreads through infected suckers and is secondarily transmitted by mealybugs in a semi-persistent manner. It causes chlorotic and necrotic streaks, bunch choking, pseudostem splitting, and in severe cases, plant dieback [8].

CMV, often referred to as Banana cucumber mosaic virus (BCMV), has a multipartite genome of three single-stranded positive-sense RNAs and a subgenomic RNA. It encodes proteins responsible for replication, movement, and encapsidation (Hayes & Buck, 1990; Palukaitis *et al*., 2003). CMV infections result in mosaic, leaf distortion, stunting, and are increasingly known as infectious chlorosis disease [9].

BBrMV, a member of the *Potyviridae* family, causes prominent reddish streaks on the pseudostem and bracts, giving rise to its name. In cultivars like Nendran and Robusta, it also induces leaf stripping and orientation changes reminiscent of a “traveler’s palm” appearance. The virus has a ∼9.7 kb positive-sense ssRNA genome coding for a large polyprotein [10].

Managing these viral infections remains a major challenge. Conventional approaches include quarantine, certification of virus-free planting materials, cultural practices such as roguing and eradication, and vector control. However, sustainable and long-term control strategies require innovative solutions, particularly those that can directly inhibit viral infection and replication [11].

One promising strategy is the molecular targeting of viral coat proteins (CPs), which play critical roles in viral infectivity, replication, and movement within host plants. The CP encapsulates the viral genome and contributes to systemic infection, symptom expression, and host specificity. Disrupting CP function through small molecules could inhibit viral life cycles and reduce disease burden [12].

In this study, we employed a structure-based virtual screening approach using molecular docking to identify potential phytochemicals with antiviral properties against the CPs of BBTV, BSV, BBrMV, and CMV. A curated library of 100 plant-derived compounds, including flavonoids, alkaloids, and terpenoids, was screened for binding affinity and interaction with viral CPs. The findings not only highlight several lead phytochemicals capable of targeting key viral proteins but also pave the way for experimental validation and eventual development of broad-spectrum, eco-friendly antiviral agents for banana disease management.

## 2. Material and Methods

### 2.1. Structural Prediction and Quality Assessment of Banana Virus Coat Proteins

The amino acid sequences of the coat proteins from four major banana viruses were retrieved from the UniProt Knowledgebase (UniProtKB). The selected proteins and their respective UniProt IDs were: *Q65386* for the capsid protein of Banana bunchy top virus (BBTV), *M1RVT2* for the genome polyprotein fragment of Banana bract mosaic virus (BBrMV), *Q8QTA0* for the capsid protein of Banana mosaic virus (used as a representative for CMV), and *A0A0K1NWL9* for the coat protein fragment of Banana streak virus (BSV). Structural models of these proteins were predicted using both SWISS-MODEL [13] and AlphaFold3 [14]. For homology modeling via SWISS-MODEL, templates were selected based on a minimum query coverage of >65% and sequence identity >30%, along with favorable Global Model Quality Estimate (GMQE) and Qualitative Model Energy Analysis (QMEAN) scores, which together ensured reliable template-targ*et al*ignments. AlphaFold3 was also used to generate high-confidence ab initio models for each protein, and all predicted structures were saved in PDB format and visualized using PyMOL v2.3.

Model quality was evaluated using the SAVES server, which included PROCHECK for generating Ramachandran plots [15], ERRAT for assessing non-bonded interactions, and Verify3D to validate residue environment compatibility. Ramachandran plots confirmed that a high percentage of residues were located within favored and allowed regions, indicating acceptable stereochemical quality. QMEAN and GMQE scores from SWISS-MODEL were used as initial metrics for model reliability, while AlphaFold3 predictions offered a complementary structure validation approach. Together, these assessments confirmed that the predicted coat protein models were structurally robust and suitable for downstream molecular docking and interaction studies.

Protein structures were prepared using the Schrödinger Maestro Protein Preparation Wizard [16]. Initial preprocessing involved optimization of hydrogen bonding networks and assignment of protonation states at physiological pH using the PROPKA [17]. Subsequently, energy minimization was performed using the OPLS4 force field [18] to achieve heavy atom convergence with a root-mean-square deviation (RMSD) threshold of 0.3 Å. Water molecules located beyond 5 Å from any ligand or active site residue were excluded from the final structure to avoid non-specific interactions [19].

### 2.2. Molecular interactions

#### 2.2.1. Ligand retrieval and preparation

A comprehensive list of phytochemicals and antiviral agents derived from *Musa acuminata*, *Musa balbisiana*, endophytic bacteria, and marine natural products (MNPs) was compiled from published literature. From this compilation, 100 representative compounds were shortlisted for molecular docking studies **(Supplementary Table S1)**. Ligand preparation was performed using the LigPrep module with the OPLS4 force field [20]. To account for physiologically relevant protonation states and stereochemical variations, Epik was employed at pH 7.0 ± 2.0 to generate tautomers and low-energy conformers, while preserving chirality. This process yielded up to 32 conformations per compound, resulting in a total of approximately 2,278 ligand structures for docking.

#### 2.2.2. Active site prediction

Active site residues were predicted using the CASTp web server [21], and the identified amino acids were selected for grid generation. Molecular docking was performed at these active sites, with a grid box centered on the predicted binding pocket **(Supplementary Table S2)**.

#### 2.2.3. Molecular Docking

Docking was restricted to ligands containing fewer than 500 atoms and no more than 100 rotatable bonds. The van der Waals radii scaling factor was set to 0.80 with a partial charge cutoff of 0.15 [22]. Flexible ligand sampling was enabled, incorporating nitrogen inversion, ring conformation sampling, and torsional adjustments favoring predicted functional groups. Intramolecular hydrogen bonding and planarity of conjugated π-systems were preserved. Docking was performed using the Extra Precision (XP) mode of the Glide module in Schrödinger. Protein–ligand interactions were visualized in 3D with PyMOL (v3.0.5) and in 2D using BIOVIA Discovery Studio Visualizer (v24.1.0.23298) [23].

### 2.3. *In silico* ADME/T and toxicity analysis

Pharmacokinetic and toxicity profiling of the top compound(s), selected based on their highest binding energies, was performed using a combination of *in silico* tools: Schrödinger’s QikProp module, SwissADME [24], ProTox-3.0 [25], ADMETlab 3.0 [26], and pkCSM [27]. These platforms provide comprehensive predictions of ADMET properties, including absorption, distribution, metabolism, excretion, toxicity, drug-likeness, and medicinal chemistry suitability of small molecules.

### 2.4. Molecular Dynamics Simulation and Trajectory Analysis

Molecular dynamics (MD) simulations of the top complexes were performed using GROMACS 2024.2 [28] with the CHARMM36 force field [29]. Systems were solvated using the TIP3P water model [30], and counterions (Na, Cl) were added for charge neutralization. After a 20,000-step energy minimization, equilibration was carried out under NVT and NPT ensembles for 100 ps each using the Nosé–Hoover thermostat and the Parrinello–Rahman barostat. A 100 ns production run was then executed.

Trajectory analyses included root mean square deviation (RMSD), root mean square fluctuation (RMSF), hydrogen bonding, radius of gyration (Rg), and solvent-accessible surface area (SASA), to evaluate complex stability, flexibility, compactness, and solvation dynamics. RMSD and Rg were also used to compute the free energy landscape (FEL), formulated as *F(X) = kBT ln P(X)*, where *P(X)* is the probability distribution along principal components. Principal component analysis (PCA) was performed by diagonalizing the covariance matrix of atomic fluctuations. Graphical representations were generated using XMGrace, VMD [31] for trajectory visualization, and Matplotlib in Python.

## 3. Results

### 3.1. Homology modelling and structural validation

Homology models of the coat proteins from Banana Bract Mosaic Virus (BBrMV), Banana Mosaic Virus (BMoV), Banana Streak Virus (BSV), and Banana Bunchy Top Virus (BBTV) were successfully generated, with predicted inter-chain confidence scores (ipTM) and overall predicted TM-scores (pTM) ranging from 0.40 to 0.75. The stereochemical integrity of all models was evaluated using the PROCHECK program, with Ramachandran plots serving as the primary validation criterion. Glycine and proline residues were excluded due to their conformational uniqueness **(Supplementary Figure S1)**.

The BBrMV model (136 residues) displayed 94.2% of residues in the most favoured regions and 5.8% in additionally allowed regions, with no residues in generously allowed or disallowed regions, indicating a high-quality structure. The BMoV model (217 residues) had 86.2% in most favored and 13.8% in allowed regions, with acceptable G-factors and no disallowed residues, supporting backbone reliability despite being slightly below the ideal threshold. The BSV model (326 residues) showed 90.6% in most favoured regions and a small fraction (0.7%) in generously allowed regions, consistent with structural flexibility and high model quality. The BBTV model (175 residues) had 88.5% in favoured, 10.9% in allowed, and 0.6% in generously allowed regions, with no disallowed residues and minimal stereochemical outliers **(Supplementary Figure S2)**.

All models exhibited acceptable G-factor values, negligible bad contacts, and >94% planarity of peptide bonds. These assessments collectively support the structural validity and suitability of the modelled proteins for downstream docking and molecular dynamics simulations **(Supplementary Table S3)**.

### 3.2. Molecular docking

Molecular docking was carried out using Schrödinger’s XP Glide protocol to explore the binding interactions between selected antiviral phytocompounds and the coat proteins of four banana-infecting viruses—Banana Bract Mosaic Virus (BBrMV), Banana Bunchy Top Virus (BBTV), Banana Mosaic Virus (BMoV), and Banana Streak Virus (BSV). The docking scores, hydrogen bonding patterns, and hydrophobic contacts were used to evaluate the stability and specificity of each protein–ligand complex **(Supplementary Table S4)**. Among the tested compounds, Saponarin consistently demonstrated the highest binding affinity and most extensive interaction profiles, suggesting its potential as a broad-spectrum antiviral agent against banana viruses **(Table 1)**.

**Table 1:** Molecular interactions and binding affinity of Saponarin based on docking scores, interacting residues and distances.

| Target | Docking score (kcal/mol) | Atoms and residues in H-bond interactions and distance (Å) | Amino acid residues in hydrophobic interactions |
| --- | --- | --- | --- |
| **BBrMV** | -13.3332 | Gly A:1 (2.90), Asp A:79 (1.96),<br>Asp A:79 (2.88), Trp A:62 (2.49) | Phe A:41 van der Waals, Ala A:45 van der Waals, Tyr A:48 van der Waals, Ile A:49 van der Waals, Tyr A:58 van der Waals, Pro A:60 van der Waals, Arg A:61 van der Waals, Thr A:2 van der Waals, Phe A:78 van der Waals, Phe A:80 van der Waals, Leu A:23 $\pi$ – alkyl, Met A:38 π-alkyl, Phe A:34 π-alkyl, Ile A:26 π-sigma, Trp A:10 π-π stacked |
| **BBTV** | -9.23 | Ser A:26 (2.63), Val A:175 (1.96), Val A:175 (1.88) | His A:174 van der Waals, Thr A:148 van der Waals, Glu A:149 van der Waals, Tyr A:62 van der Waals, Arg A:61 van der Waals, Ala A:29 van der Waals, Lys A:27 van der Waals, Lys A:145 van der Waals, Phe A:63 van der Waals, Trp A:65 π-π stacked (5.17), |
| **BMoV** | -8.75 | Lys A:65 (1.70), Lys A:65 (183), Asp A:169 (2.12), Asp A:169 (2.42), Arg A:106 (2.44), Glu A:62 (1.86), Glu A:62 (2.79), Arg A:63 (2.12), Arg A:63 (2.28), Arg A:63 (2.77), Lys A:44 (1.99) | Lys A:65 π-Cation (to aromatic ring) (4.80, 4.82), His A:174 van der Waals, Ser A:171 van der Waals, Ser A:105 van der Waals, Leu A:167 van der Waals, Pro A:66 van der Waals, Cys A:64 van der Waals |
| **BSV** | -11.7029 | Asn A:50 (1.75), Asn A:50 (2.90), Asn A:50 (2.02), Glu A:246 (2.30), Asp A:249 (2.47) | Arg A:47 van der Waals, Asp A:42 van der Waals, Met A:247 van der Waals, Thr A:43 van der Waals, Phe A:250 van der Waals, Lys A:251 van der Waals, Tyr A:46 van der Waals, Asn A:252 van der Waals, Lys A:107 van der Waals, Tyr A:253 π-Alkyl (ligand ring→Tyr) (4.93) |

Saponarin exhibited a GlideScore of –13.33 kcal/mol against the BBrMV coat protein, the strongest among all ligand–target combinations. It formed multiple strong hydrogen bonds with Gly1, Asp79 (two interactions), and Trp62 at distances ranging from 1.96 to 2.90 Å. Extensive hydrophobic interactions involved 17 residues, including key π-alkyl contacts with Phe34, Met38, Ile26, and π–π stacking with Trp10, further stabilizing the complex. These results reflect high specificity and binding strength within the predicted active pocket **(Figure 1)**.

**Figure 1:**
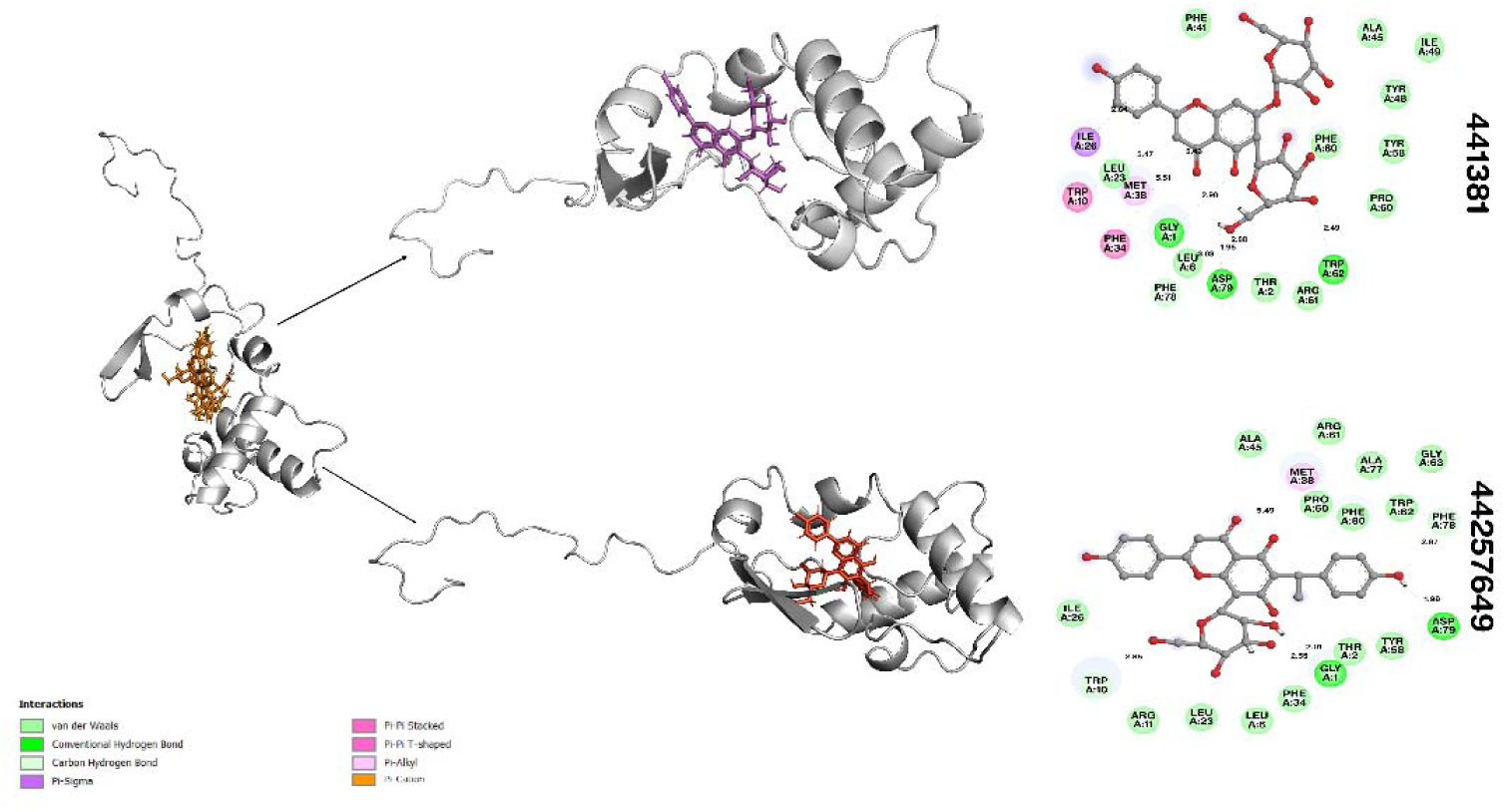
Molecular docking poses and 2D interaction diagrams for top-scoring compounds with the Banana bract mosaic virus (BBrMV) coat protein. Left: Cartoon representations of the BBrMV coat protein with docked compounds (PubChem IDs: 441381 [top, purple], 44257649 [bottom, orange]). Middle: Enlarged views showing the binding orientation of each compound within the active site pocket. Right: Two-dimensional interaction diagrams highlighting the key amino acid residues involved in ligand binding for each compound, as predicted by docking analysis. Hydrogen bonds and hydrophobic interactions are indicated, with bond distances labelled in angstroms. These visualizations demonstrate the specific interactions mediating compound binding to the viral coat protein.

**Figure 2:**
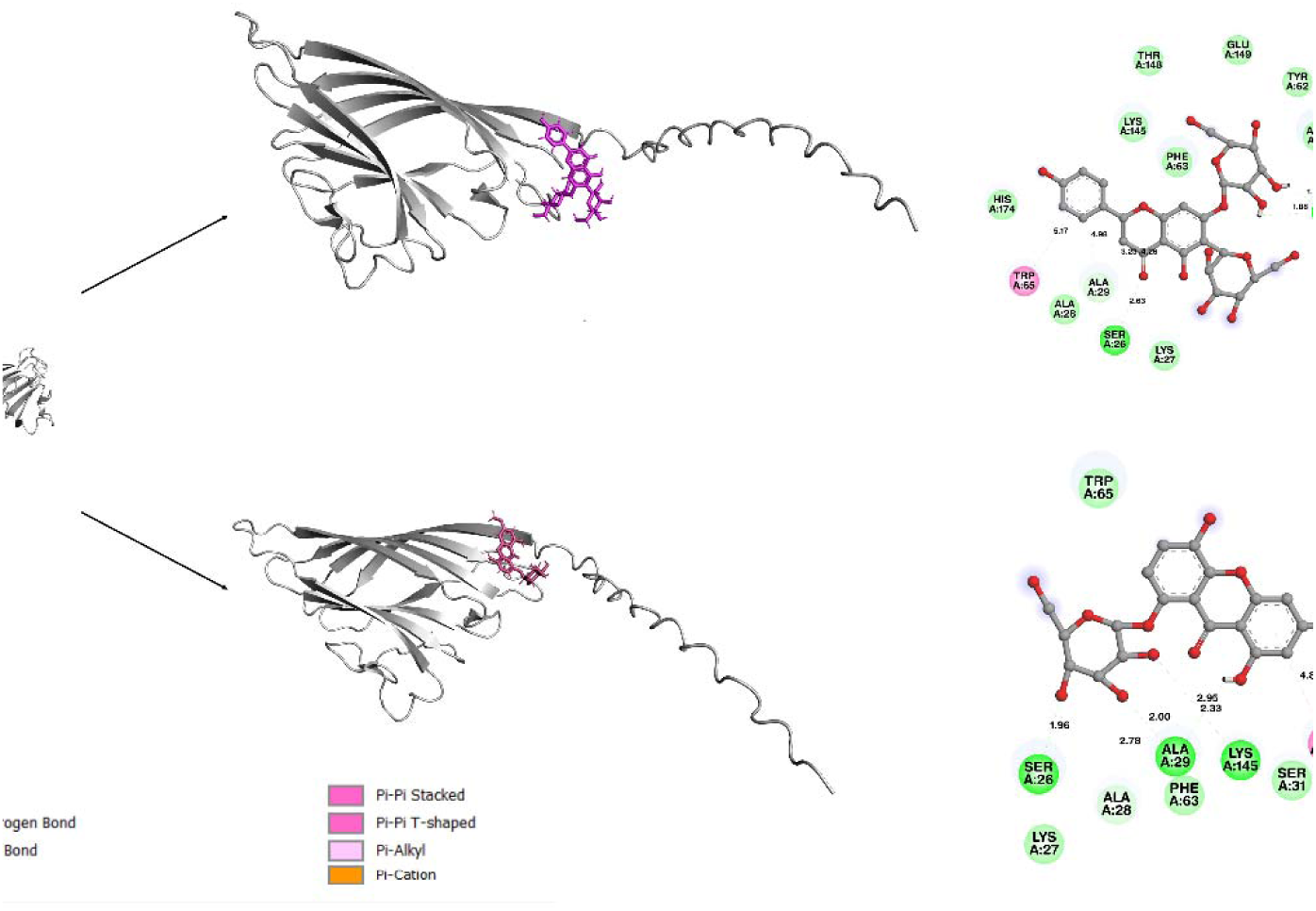
Molecular docking poses and 2D interaction diagrams for top-scoring compounds with the Banana bract mosaic virus (BBTV) coat protein.]). Left: Cartoon representations of the BBTV coat protein with docked compounds (PubChem IDs: 441381 [top, purple], 5858086 [bottom, orange]). Middle: Enlarged views showing the binding orientation of each compound within the active site pocket. Right: Two-dimensional interaction diagrams depicting the interactions between each ligand and the surrounding amino acid residues, as identified in the docking analysis. Hydrogen bonds and hydrophobic interactions are highlighted, with bond distances indicated in angstroms. These visualizations illustrate the molecular basis for ligand binding and support the selection of promising antiviral candidates.

Against the BBTV coat protein, Saponarin recorded a docking score of –9.23 kcal/mol, forming three hydrogen bonds with Ser26 and Val175, with close contact distances (1.88– 2.63 Å). Eleven residues were involved in hydrophobic interactions, including π–π stacking with Trp65 (5.17 Å) and van der Waals contacts with critical interface residues such as Tyr62, Arg61, and Phe63. These interactions support the stable occupancy of Saponarin in the binding site of BBTV **(Figure 1)**.

Saponarin displayed strong binding to the BMoV coat protein, with a GlideScore of – 8.75 kcal/mol. Notably, it engaged in an extensive hydrogen bonding network with Lys65 (two bonds), Asp169 (two bonds), Arg106, Glu62 (two bonds), Arg63 (three bonds), and Lys44, totaling 11 hydrogen bonds, indicating high interaction specificity. Additionally, π–cation interactions with Lys65 and van der Waals contacts with residues such as His174, Pro66, and Cys64 enhanced the complex stability **(Figure 3)**.

**Figure 3:**
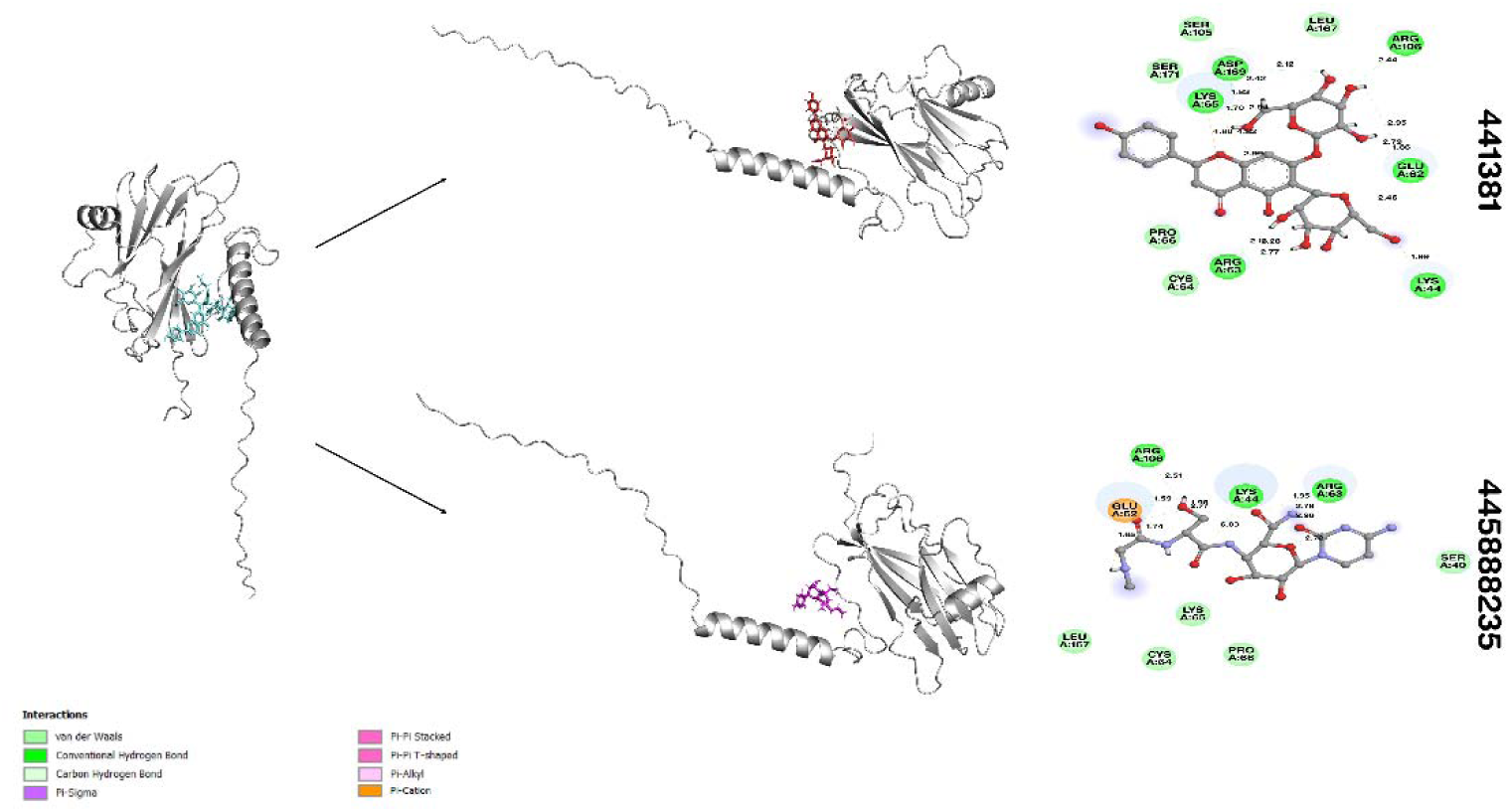
Molecular docking poses and 2D interaction diagrams for top-scoring compounds with the Banana mosaic virus (BMoV) coat protein. Left: Cartoon representations of the BMoV coat protein with docked compounds (PubChem IDs: 441381 [top, red], 44588235 [bottom, magenta]). Middle: Enlarged views illustrating the binding orientation of each compound within the active site pocket. Right: Two-dimensional interaction diagrams showing key interactions between each ligand and neighbouring amino acid residues, as determined by docking analysis. Hydrogen bonds, hydrophobic contacts, and bond distances (in angstroms) are annotated. These results highlight the specific molecular interactions responsible for strong binding affinities of these antiviral candidates.

Among all compounds tested against the BSV coat protein, Saponarin again showed the best binding affinity, with a Glide Score of –11.70 kcal/mol. It formed five hydrogen bonds with Asn50 (three interactions), Glu246, and Asp249, and established hydrophobic interactions with key residues including Tyr46, Met247, and Phe250. A notable π–alkyl interaction with Tyr253 further reinforced ligand anchoring. This interaction profile highlights strong molecular complementarity between Saponarin and the BSV protein binding pocket **(Figure 4)**.

**Figure 4:**
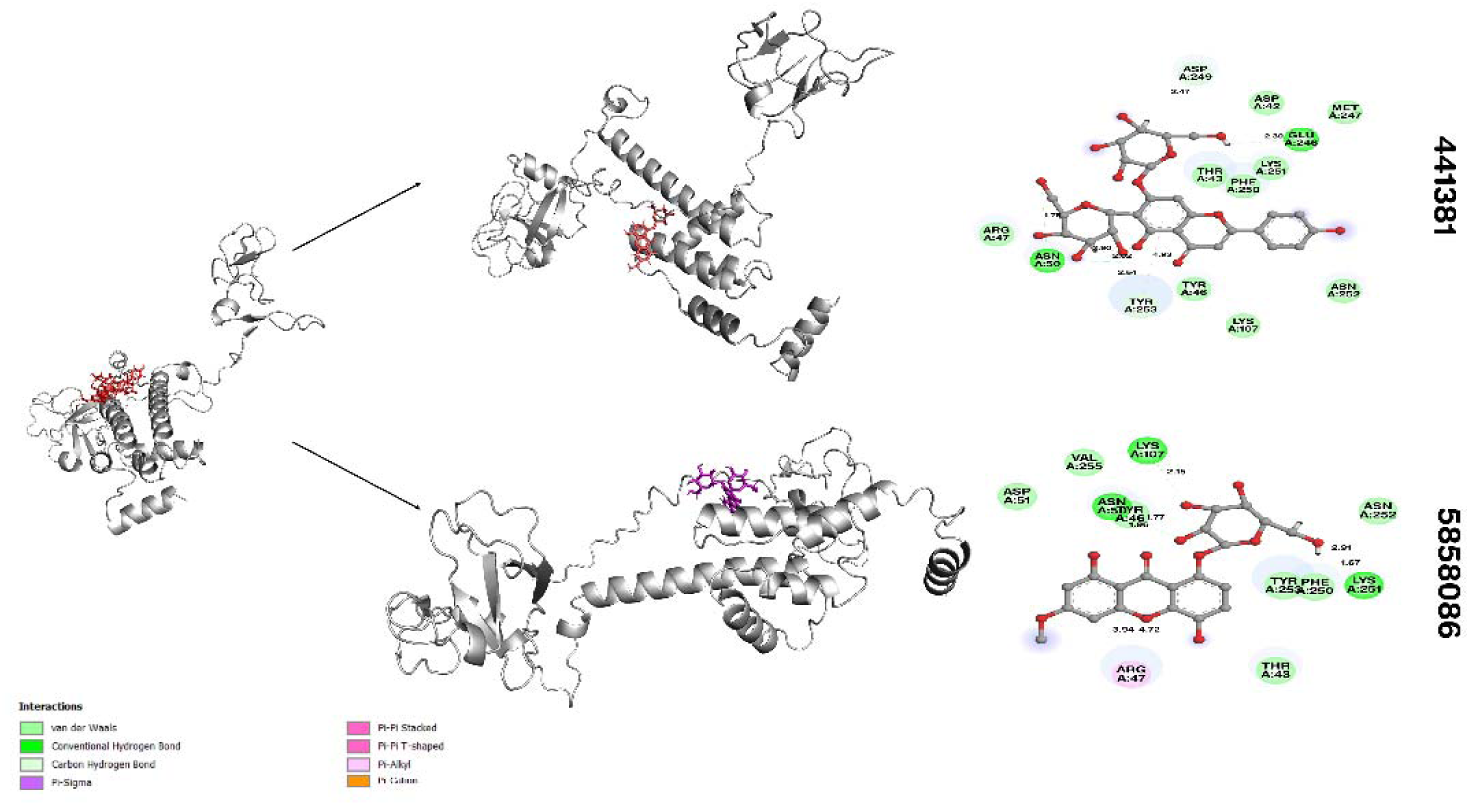
Molecular docking poses and 2D interaction diagrams for top-scoring compounds with the Banana streak virus (BSV) coat protein. Left: Cartoon representations of the BSV coat protein with docked compounds (PubChem IDs: 441381 [top, red], 5858086 [bottom, magenta]). Middle: Enlarged structural views highlighting the binding orientation of each compound within the predicted active site pocket. Right: Two-dimensional interaction diagrams illustrating key interactions between each ligand and surrounding amino acid residues, as identified in the docking analysis. Hydrogen bonds, hydrophobic contacts, and corresponding bond distances (in angstroms) are annotated. These results demonstrate the molecular determinants of binding for the most promising antiviral candidates targeting BSV.

Compared to other tested ligands, Saponarin consistently yielded the most favorable docking scores across all viral targets, ranging from –8.75 to –13.33 kcal/mol, and exhibited the most comprehensive hydrogen bonding and hydrophobic interaction profiles. Its ability to form multiple, close-contact hydrogen bonds and stabilize through π-based interactions suggests strong target engagement, reinforcing its role as the top-scoring candidate for further experimental validation.

### 3.3. Pharmacokinetic Profiling of Saponarin

Saponarin, a flavonoid glucoside with a molecular weight of 594.52 Da and polar surface area of 235.78 Å², exhibits physicochemical properties highly favorable for interaction with polar surface residues of viral coat proteins from banana-infecting viruses such as BBTV, BBrMV, BSV, and BMoV **(Figure 5)**. Its high number of hydrogen bond acceptors (15) and donors (10), combined with six rotatable bonds, enables strong, stable, and flexible binding within the structurally conserved groove and capsid interface pockets of the coat proteins, a supported by molecular docking and MD data. The compound’s hydrophilic nature (logP = - 2.4352) supports xylem transport and aqueous formulation, ensuring effective translocation through plant vasculature and sustained bioavailability at the site of infection. Despite moderat water solubility (logS = -2.724), its low systemic clearance (log CL = -0.047) and absence of rapid efflux properties favor its persistence within plant tissues, increasing the duration of antiviral activity. Environmentally, Saponarin poses minimal ecotoxicological risks, with extremely low aquatic organism toxicity (minnow log mM = 11.323, *T. pyriformis* toxicity = 0.285 µg/L), and its non-mutagenic, non-sensitizing profile ensures safety in agroecosystems. As a non-inhibitor of major metabolic enzymes and transporters, and with no hepatotoxic or skin- sensitizing effects, Saponarin is suitable for integration into sustainable plant protection strategies (**Supplementary Table S5)**.

**Figure 5:**
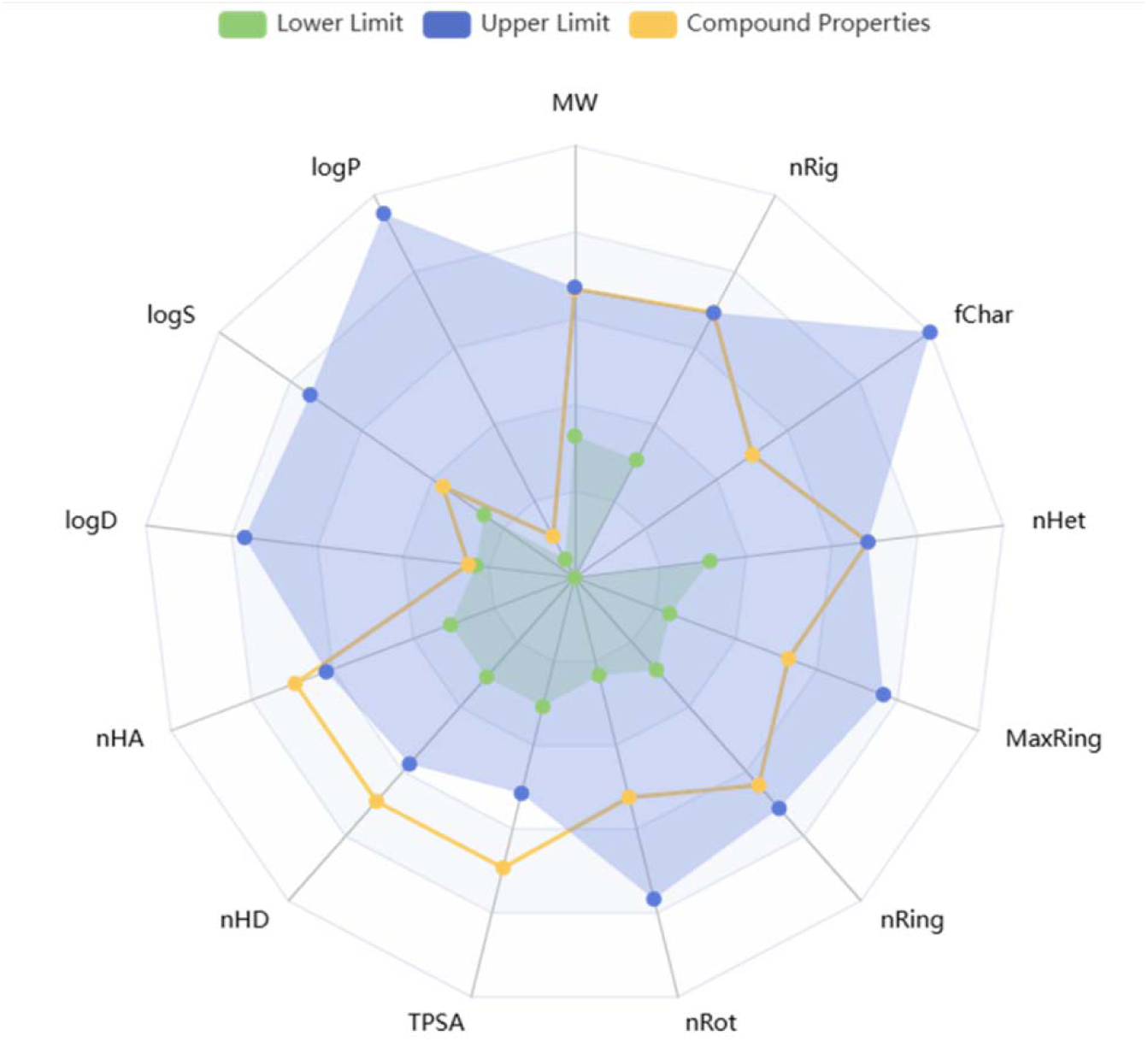
Bioavailability radar plot of Saponarin obtained from AdmetLab 3.0.

### 3.4. Molecular dynamics simulation

To investigate the dynamic stability and conformational behavior of the top compound Saponarin bound to four banana viral coat proteins—BBrMV, BBTV, BMoV, and BSV—100-ns all-atom molecular dynamics (MD) simulations were conducted. The simulations were evaluated using several parameters, including protein and ligand RMSD, RMSF, radius of gyration (Rg), hydrogen bond occupancy, and solvent-accessible surface area (SASA).

#### 3.4.1. Root Mean Square Deviation (RMSD)

##### 3.4.1.1. RMSD of Protein Backbone

The backbone RMSD profiles revealed that all four protein–ligand complexes attained equilibrium within the initial 10–15 ns and remained stable throughout the 100-ns simulation. In the BBrMV–Saponarin complex, RMSD increased from ∼1 Å to 2.3 Å within the first 10 ns, stabilizing thereafter at 2.2 ± 0.2 Å. The BBTV–Saponarin complex reached equilibrium at 2.5 Å by 15 ns and remained consistent around 2.4 ± 0.25 Å. Similarly, the BMoV–Saponarin complex stabilized at ∼2.2 Å after 12 ns, maintaining minimal fluctuations (2.1 ± 0.15 Å). The BSV–Saponarin complex showed slightly more variation but still achieved stable RMSD values between 2.3–2.5 Å (mean 2.4 ± 0.2 Å). These results confirm that Saponarin binding imparts structural integrity to all target proteins over extended timescales **(Panel A of Figure 6**, **Figure 7**, **Figure 8**, **Figure 9)**.

**Figure 6:**
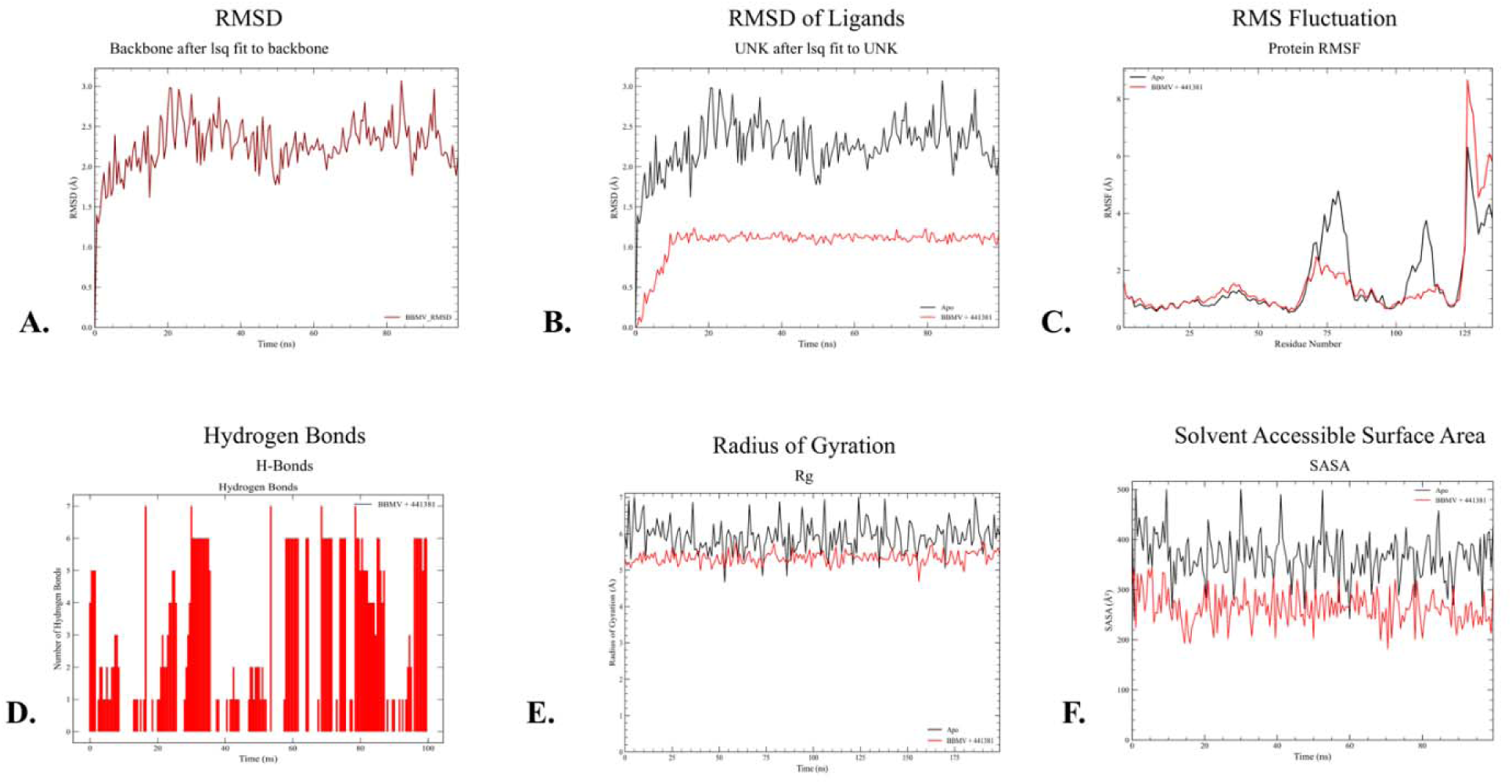
Comparative molecular dynamics analysis of BBrMV in apo form (black) and in complex with Saponarin (red). (A) Backbone RMSD indicating structural stability of the complex. (B) Ligand RMSD showing consistent binding of Saponarin. (C) RMSF depicting reduced residue flexibility upon ligand binding. (D) Number of hydrogen bonds between BBrMV and Saponarin over time. (E) Radius of gyration (Rg) illustrating compactness of the complex. (F) Solvent accessible surface area (SASA) reflecting reduced solvent exposure upon complex formation.

**Figure 7:**
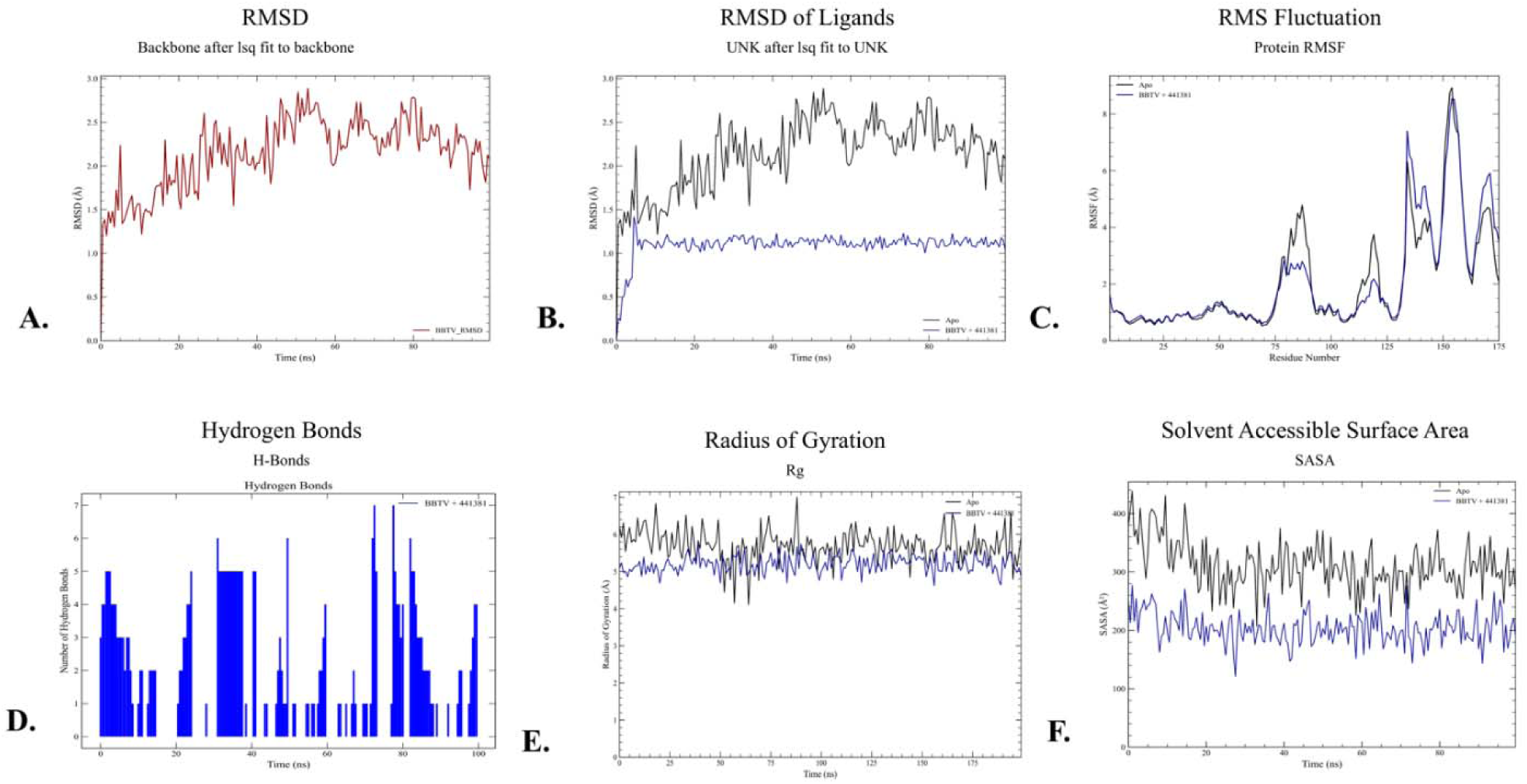
Comparative molecular dynamics analysis of BBSV in apo form (black) and in complex with Saponari (red). (A) Backbone RMSD indicating structural stability of the complex. (B) Ligand RMSD showing consistent binding of Saponarin. (C) RMSF depicting reduced residue flexibility upon ligand binding. (D) Number of hydrogen bonds between BBSV and Saponarin over time. (E) Radius of gyration (Rg) illustrating compactness of the complex. (F) Solvent accessible surface area (SASA) reflecting reduced solvent exposure upon complex formation.

**Figure 8:**
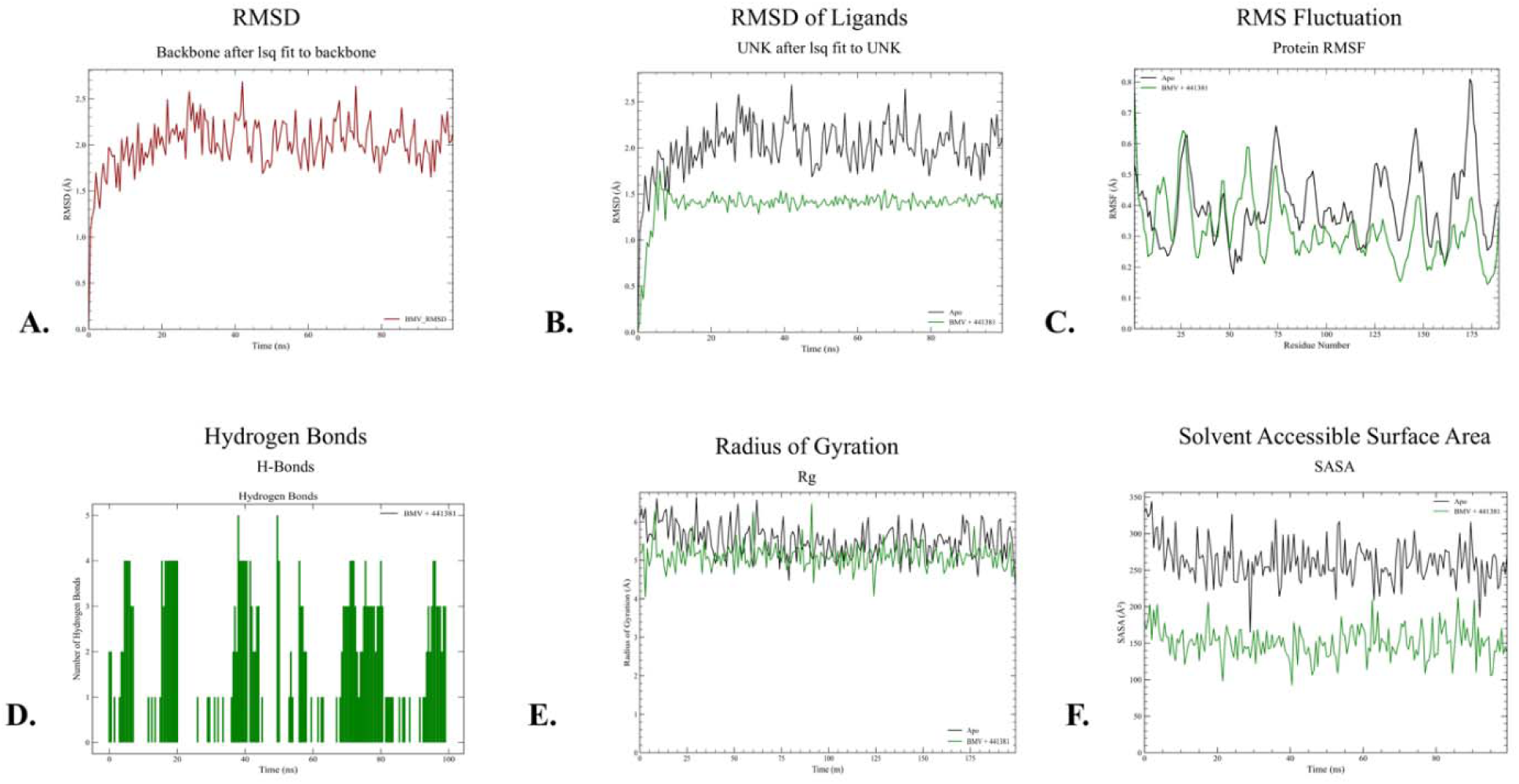
Comparative molecular dynamics analysis of BMoV in apo form (black) and in complex with Saponarin (red). (A) Backbone RMSD indicating structural stability of the complex. (B) Ligand RMSD showing consistent binding of Saponarin. (C) RMSF depicting reduced residue flexibility upon ligand binding. (D) Number of hydrogen bonds between BMoV and Saponarin over time. (E) Radius of gyration (Rg) illustrating compactness of the complex. (F) Solvent accessible surface area (SASA) reflecting reduced solvent exposure upon complex formation.

**Figure 9:**
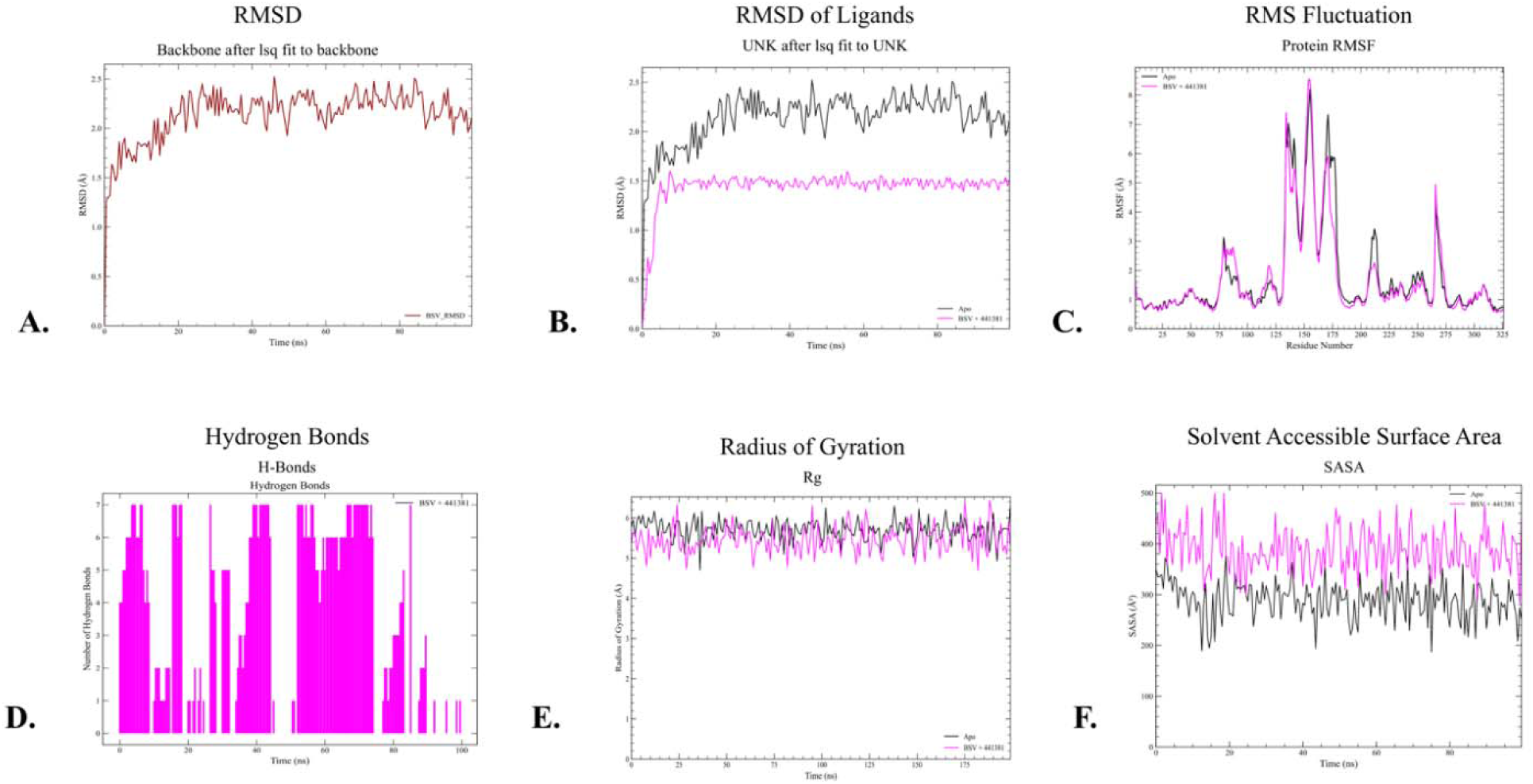
Comparative molecular dynamics analysis of BSV in apo form (black) and in complex with Saponarin (red). (A) Backbone RMSD indicating structural stability of the complex. (B) Ligand RMSD showing consistent binding of Saponarin. (C) RMSF depicting reduced residue flexibility upon ligand binding. (D) Number of hydrogen bonds between BSV and Saponarin over time. (E) Radius of gyration (Rg) illustrating compactness of the complex. (F) Solvent accessible surface area (SASA) reflecting reduced solvent exposure upon complex formation.

##### 3.4.1.2. RMSD of Saponarin

Ligand RMSD analyses confirmed high binding pose stability across all systems. In BBrMV and BMoV, Saponarin’s RMSD remained within 1.5–2.3 Å, suggesting tight retention in the binding pocket. For BBTV and BSV, the ligand RMSD values fluctuated between 1.8–2.5 Å with occasional transient spikes up to ∼3 Å, quickly returning to baseline—indicative of minor conformational adjustments rather than unbinding. Overall, Saponarin maintained RMSD <2.5 Å for >90% of the simulation time across all systems, affirming pose fidelity and stable binding **(Panel B of Figure 6**, **Figure 7**, **Figure 8**, **Figure 9)**.

#### 3.4.2. Root Mean Square Fluctuation (RMSF)

Residue-level flexibility was assessed using RMSF analysis. Across all four coat proteins, backbone fluctuations were largely confined to loop regions—typically residues 20–60 and 130– 160—while secondary structure elements remained stable (RMSF <1.5 Å). Binding pocket residues involved in hydrogen bonding and hydrophobic interactions with Saponarin (e.g., Trp10, Arg26, Phe34 in BBrMV; Lys23, Arg144 in BBTV; Gln37, Arg33 in BMoV; Thr41, Tyr46 in BSV) consistently showed RMSF values <1.8 Å. These observations highlight localized flexibility while preserving a stabilized binding environment for the ligand. The overall mean RMSF across all proteins was 1.65 ± 0.3 Å **(Panel C of Figure 6**, **Figure 7**, **Figure 8**, **Figure 9)**.

#### 3.4.3. Radius of Gyration (Rg)

The radius of gyration (Rg) analysis provided insights into the global compactness of the protein structures. All complexes achieved stable Rg values around 19 Å, with minimal fluctuation (<±0.25 Å) after the initial 20 ns. BBTV and BSV complexes converged more rapidly (∼15 ns) and exhibited the tightest Rg distributions, indicating restrained conformational flexibility. Slightly higher Rg variance (±0.3–0.4 Å) in BBrMV and BMoV reflected minor mobility in loop regions, without compromising structural integrity. Overall, these results suggest sustained compactness and well-folded structures across all complexes **(Panel D of Figure 6**, **Figure 7**, **Figure 8**, **Figure 9)**.

#### 3.4.4. Hydrogen Bond Analysis

Saponarin formed persistent hydrogen bonds with key residues in all complexes, contributing to binding stability. In the BBrMV complex, strong and consistent interactions were observed with Trp10 (90% occupancy) and Arg26. The BBTV complex showed hydrogen bonds with Lys23 (87%) and Arg144 (85%). For BMoV, sustained interactions were seen with Gln37 (80%) and Arg33 (78%). In the BSV complex, Thr41 and Tyr46 maintained H-bond occupancies of 83% and 80%, respectively. Notably, no prolonged periods (>15 ns) of H-bond absence were recorded in any system, indicating continuous ligand engagement **(Panel E of Figure 6**, **Figure 7**, **Figure 8**, **Figure 9)**.

#### 3.4.5. Solvent Accessible Surface Area (SASA)

SASA analysis was conducted to assess the burial of Saponarin within the protein binding sites and solvent exposure of the complexes. Across all systems, SASA values started between 150–200 Å² and converged to 160–180 Å² by ∼30 ns, maintaining stability (±40 Å²) for the remainder of the simulation. BBTV and BSV showed narrower SASA fluctuation range (±30 Å²), suggesting tighter ligand integration into the binding pocket. Slightly wider variations in BBrMV and BMoV (±50 Å²) indicate flexible but non-disruptive pocket dynamics. Overall, the SASA results reinforce the consistent burial of Saponarin in the receptor cavities **(Panel F of Figure 6**, **Figure 7**, **Figure 8**, **Figure 9)**.

## 4. Discussion

Our homology-modelled coat proteins of BBrMV, BMoV, BSV, and BBTV exhibited high stereochemical quality (up to 94% residues in most-favoured regions, negligible outliers), establishing a reliable structural framework for subsequent docking and dynamics analyses [32]. The exceptional docking score achieved by Saponarin across all targets (–13.33 to – 8.75 kcal/mol) indicates strong and specific complementarity with coat protein pockets, further corroborated by robust hydrogen bonding and π-based hydrophobic contacts. Notably, key aromatic residues—Trp10/Trp62 in BBrMV, Trp65/Tyr62 in BBTV, His174/Arg106 in BMoV, and Tyr46 in BSV—mediated π–π stacking and π–alkyl interactions, consistent with other flavonoid–virus docking studies illustrating the critical role of aromatic residues at capsid interfaces [33–36]. The concentration of these interactions around conserved intersubunit grooves suggests an allosteric mode of inhibition—binding outside the canonical assembly interfaces, yet likely disrupting capsid stabilization or inter-protein contacts critical to viral infectivity [37,38]. This is supported by MD simulations, where sustained RMSD (∼2.1–2.5 Å), stable Rg (∼19 Å), high hydrogen bond occupancies (>80%), and steady SASA profiles over 100 ns confirm that Saponarin remains tightly anchored, preserving structural integrity rather than causing destabilization—hallmarks of allosteric modulators [39]. As a flavonoid glucoside, Saponarin’s physicochemical properties (15 H-bond donors/acceptors, logP = –2.43, logS = – 2.72, PSA = 235 Å²) facilitate aqueous solubility and polar recognition, corroborating its proposed xylem translocation in planta, paralleling the behavior reported for saponin/flavonoid transport in plants . Moreover, its favorable pharmacokinetic profile—low clearance, absence of mutagenicity or hepatotoxicity—mirrors other structurally-similar phytocompounds with broad-spectrum antiviral potential, while ecotoxicity assessments affirm its environmental safety. Combined, these data present compelling evidence that Saponarin could serve as a biofriendly antiviral agent in banana cultivation systems. Nonetheless, limitations remain - our findings derive entirely from *in silico* modeling, without corroborative *in vitro* binding, capsid assembly, or plant infectivity assays. Future work should include biophysical characterization (e.g., SPR, ITC), viral infectivity assays under greenhouse conditions, and truncated protein or pseudo-capsid binding studies to validate assembly inhibition [40]. Additionally, exploring synergy with other banana-derived saponins or comparison with known antiviral lectins like Banana lectin (BanLec) [41] would strengthen functional relevance. Scalability of Saponarin production—via extraction from barley sprouts, barley leaves, or grape leaves—and formulation into foliar sprays or soil drenches should also be evaluated in field trials to assess stability, dosage, and systemic distribution [42–44]. Leveraging its physicochemical and ecotoxicological profile, Saponarin thus holds promise as a sustainable, allosteric inhibitor of banana viral coat proteins—but bridging computational and empirical evidence remains the critical next step toward deployment in integrated virus management strategies.

## 5. Conclusion

This study identifies Saponarin as a promising broad-spectrum antiviral phytocompound targeting the coat proteins of four major banana-infecting viruses—BBrMV, BBTV, BMoV, and BSV—through structure-based molecular docking and all-atom MD simulations. Its high binding affinity, extensive hydrogen bonding, and stable interaction with conserved capsid interface residues across diverse viral targets suggest a potential allosteric mechanism of inhibition that could interfere with capsid assembly or stability. The compound’s favorable physicochemical and pharmacokinetic properties, combined with its minimal ecological toxicity, further support its applicability as a safe and effective agent for plant virus management. These findings provide a rational basis for advancing Saponarin toward experimental validation and formulation for sustainable agricultural deployment against banana viral diseases.

## 6. Declaration

### 6.2. Authorship Contribution Statement

**Mohammad Imran Kumasagi**: Conceptualization, Methodology, Formal analysis, Investigation, Writing – original draft, Visualization. **Nagesha N**: Supervision, Project administration, Conceptualization. **Dyumn Dwivedi**: Formal analysis, Investigation, Writing – original draft, Writing – review & editing, Visualization. **Shankarappa K.S.**: Supervision, Project administration. **Uddalak Das**: Supervision, Writing – review & editing, Funding acquisition.

## Acknowledgement

The authors would like to thank the Department of Plant Biotechnology, College of Agriculture, University of Agricultural Sciences, Bangalore and the School of Biotechnology, Jawaharlal Nehru University, for providing the facilities and also for their constant support in carrying out the research work. We would like to express their sincere gratitude to Prof. R. Sowdhamini from the National Centre for Biological Sciences, TIFR, for generously providing the Schödinger drug discovery suite, which was instrumental in carrying out the drug discovery and screening operations. We also extend our heartfelt thanks to Mr. Amey Ghodeshwar, NomadX Holdings LLC, for his invaluable assistance in providing system access and for offering the simulation suite that enabled the successful execution of our simulations.

## 6.4. Ethical Statements

None

## 6.5. Declaration of generative AI and AI-assisted technologies in the writing process

The writing of this research paper involved the use of generative AI and AI-assisted technologies only to enhance the clarity, coherence, and overall quality of the manuscript. The authors acknowledge the contributions of AI in the writing process only. All interpretations and conclusions drawn in this manuscript are the sole responsibility of the author.

