## Supplementary File for "Structure-based discovery of Saponarin as a broad-spectrum allosteric inhibitor of banana viral coat proteins"

**Supplementary Table S1:** List of ligands used in molecular docking against four viral coat proteins of banana viruses

| Group | Compound | Pubchem ID | Source |
| --- | --- | --- | --- |
| Alkaloid | Trigonelline | 5570 | *Hypoestesphyllostachya* |
| Antioxidants | Beta ionone | 638014 | *Cedrus* |
|  | Cucumerin-A | 44257649 | *Cucumis sativus* L |
|  | Cucumerin-B | 44257648 | *Cucumis sativus* L |
|  | 4-Hydroxycinnamic acid | 637542 | *Musa buibisiana* |
| Antiviral agents | Carageenan | 71597331 | *Acanthophoraspecifira* |
|  | Acyclovir | 135398513 | Chemically synthesised |
|  | 5-Azacytidine | 9444 | Chemically synthesised |
|  | Cytarabine | 6253 | Chemically synthesised |
|  | Ribavirin | 37542 | Chemically synthesised |
|  | Ningnanmycin | 44588235 | *Streptomyces noursei* |
|  | Vidarabine | 21704 | Chemically synthesised |
|  | Acycloguanosine | 135398513 | Chemically synthesised |
|  | 2-Thiouracil | 1269845 | Chemically synthesised |
|  | Moroxydine hydrochloride | 76621 | Chemically synthesised |
|  | Luotonin A | 10334120 | *Peganumnigella strum* |
|  | Pergularinine | 264751 | *Cynanchum, Pergularia,* |
|  | Antofine | 639288 | *Cynanchumkomarovii* |
|  | Deoxytylophorinine | 6426880 | *Cynanchumkomarovii* |
|  | Pyrroloisoquinoline | 86733878 | *Cynanchumkomarovii* |
|  | Streptindole | 135431 | *Streptococcus faecium* |
|  | Tryptanthrin | 73549 | *Indigofera tinctoria* L |
|  | Essramycin | 24829329 | *Streptomyces sp* |
|  | Chlorogenic acid | 1794427 | *Solanum tuberosum* L |
|  | Peonidin | 441773 | *Solanum tuberosum* L |
|  | Swertianolin | 5858086 | *Swertia chirayita*L*,* |
|  | Zidovudine | 35370 | Chemically synthesised |
| Carotenoid | Lutein | 5281243 | *Cucurbita pepo* |
| Flavonoids | Isoscoparin | 442611 | *Cucumis sativus* L |
|  | Saponin | 441381 | *Cucumis sativus* L |
|  | Vicenin-2 | 442664 | *Cucumis sativus* L |
|  | Apigenin -7-o-glucoside | 5280746 | *Cucumis sativus* L |
|  | Quercetin -3-o-glucoside | 5280804 | *Cucumis sativus* L |
|  | Isorhamnetin -3-o-glucoside | 5318645 | *Cucumis sativus* L |
|  | Kaemferol -3-o-rhamnoside | 5316673 | *Cucumis sativus* L |
|  | Kaempferol | 5280863 | *Musa acuminata* |
|  | Myricetin | 5281672 | *Musa acuminata* |
|  | Quercetin | 5280343 | *Musa acuminata* |
|  | (-)-Epigallocatechin gallate | 65064 | *Musa acuminata* |
|  | Apigenin | 5280443 | *Musa acuminata* |
|  | Luteolin | 5280445 | *Musa acuminata* |
|  | (-)-Epicatechin | 72276 | *Musa acuminata* |
|  | (+)-Gallocatechin | 65084 | *Musa acuminata* |
|  | Gallocatechin gallate | 5276890 | *Musa acuminata* |
|  | (-)-Epicatechin gallate | 107905 | *Musa acuminata* |
|  | Epigallocatechin | 72277 | *Musa acuminata* |
|  | Procyanidin C1 | 169853 | *Musa acuminata* |
|  | Procyanidin B4 | 147299 | *Musa acuminata* |
|  | Procyanidin B1 | 11250133 | *Musa acuminata* |
|  | Procyanidin B8 | 474541 | *Musa acuminata* |
|  | Procyanidin B3 | 146798 | *Musa acuminata* |
|  | Procyanidin B2 | 122738 | *Musa acuminata* |
|  | Prodelphinidin B | 5089687 | *Musa acuminata* |
|  | Vitexin | 5280441 | *Cucumis sativus* L |
|  | Isovitexin | 162350 | *Cucumis sativus* L |
|  | Orientin | 5281675 | *Cucumis sativus* L |
|  | Isoorientin | 114776 | *Cucumis sativus* L |
| lignans | Pinoresinol | 73399 | *Musa acuminata* |
|  | Matairesinol | 119205 | *Musa acuminata* |
|  | Lariciresinol | 332427 | *Musa acuminata* |
|  | Secoisolariciresinol | 65373 | *Musa acuminata* |
| Phenols | 4'-O-Methylirenolone | 638391 | *Musa acuminata* |
|  | Cucurbitoside K- | 11656306 | *Cucurbita pepo* |
|  | Cucurbitoside J | 11497614 | *Cucurbita pepo* |
|  | Cucurbitoside M | 11519464 | *Cucurbita pepo* |
|  | Cucurbitoside F | 11555374 | *Cucurbita pepo* |
|  | Cucurbitoside H | 11512173 | *Cucurbita pepo* |
|  | Cucurbitoside I | 11555787 | *Cucurbita pepo* |
|  | Cianidanol | 9064 | *Musa acuminata* |
|  | Gallic acid | 370 | *Musa acuminata* |
|  | Bryononic acid | 472768 | *Benincasa hispida* |
|  | p-Coumaraldehyde | 641301 | *Aralia bipinnata* |
| Phytoalexins | Irenolone | 2754650 | *Musa acuminata* |
|  | Hydroxyanigorufone | 11471752 | *Musa acuminata* |
|  | Anigorufone | 636472 | *Musa acuminata* |
|  | Musanolone F | 10734307 | *Musa acuminata* |
|  | Musanolone E | 10733280 | *Musa acuminata* |
| Taninns | (-)-Musellarin A | 122380090 | *Musa acuminata* |
|  | Musellarin A | 85413490 | *Musa acuminata* |
| Terpinoids | Musabalbisiane B | 131752423 | *Musa buibisiana* |
|  | (2R,3R)-2,3-bis[(4-hydroxy-3-methoxyphenyl)(113C)methyl](113C)butane-1,4-diol | 11646293 | *Musa acuminata* |
|  | (2R,3R)-2-(3,4-dihydroxyphenyl)-6-[(2R,3S,4S)-2-(3,4-dihydroxyphenyl)-3,5,7-trihydroxy-3,4-dihydro-2H-chromen-4-yl]-3,4-dihydro-2H-chromene-3,5,7-triol | 11649946 | *Musa acuminata* |
|  | Musabalbisiane A | 131752422 | *Musa buibisiana* |
|  | Musabalbisiane C | 131752424 | *Musa buibisiana* |
|  | 2-[(1R,2R,2'R,4aR,5R,5'S,6R,8R,8aS)-5'-(furan-3-yl)-2',8-dihydroxy-1,4a,6,8atetrakis(hydroxymethyl)-2-[(Z)-2-methylbut-2-enoyl]oxyspiro[2,3,4,6,7,8hexahydronaphthalene-5,3'-oxolane]-1-yl]acetic acid | 163190920 | *Musa buibisiana* |
|  | 1,7-Bis(4-hydroxyphenyl)hepta-4,6-dien-3-one | 10613719 | *Musa buibisiana* |
|  | (4E,6E)-1,7-bis(4-hydroxyphenyl)-4,6-heptadien-one | 71447176 | *Musa buibisiana* |
|  | 2,3-Dihydro-4-(4-methoxyphenyl)-1H-phenalene-1alpha,2beta,3beta-triol | 10424295 | *Musa buibisiana* |
|  | 2-(4-Hydroxyphenyl)naphthalic anhydride | 10613719 | *Musa buibisiana* |
|  | Cucurbitacin-A | 5281315 | *Cucumis sativus* L |
|  | Cucurbitacin-B | 5281316 | *Cucumis sativus* L |
|  | Cucurbitacin-C | 5281317 | *Cucumis sativus* L |
|  | Cucurbitacin-D | 5281318 | *Cucumis sativus* L |
|  | Cucurbitacin-E | 5281319 | *Cucumis sativus* L |
|  | Cucurbitacin-I | 5281321 | *Cucumis sativus* L |
|  | Indole-3-aldehyde | 10256 | *Euphorbia hirsuta L* |
|  | Indole-3-carboxylic acid | 69867 | *Cucumis sativus* L |
|  | (+)-Dehydrovomifoliol | 688492 | *Cucumis sativus* L |
|  | Cucumegastigmane-I | 16105430 | *Cucumis sativus* L |
|  | Cucumegastigmane-II | 16105434 | *Cucumis sativus* L |

**Supplementary Table S2.** Detailed active site characteristics of modelled banana viral coat proteins as predicted by CASTp.

| Virus | Active Site Residues | Surface Area (Å²) | Pocket Volume (Å³) |
| --- | --- | --- | --- |
| Banana bract mosaic virus (BBMV) | Gly1, Thr2, Leu6, Thr9, Trp10, Arg11, Met13, Leu23, Ile26, Phe34, Met38 | 499.1 | 958.3 |
| Banana bunchy top virus (BBTV) | Arg21, Lys23, Tyr24, Ser26, Ala28, Phe63, Trp65, Arg144, Lys145 | 141.7 | 483.8 |
| Banana mosaic virus (BMV) | Arg22, Arg33, Gln37, Ser40, Arg41, Lys44, Ala47, Ala48, Gly49, Gly50 | 705.9 | 2492.8 |
| Banana streak virus (BSV) | Met40, Thr41, Asp42, Ile45, Tyr46, Met49, Asn50, Phe53, Trp63, Leu80, Phe88 | 1977.3 | 2960.5 |

**Supplementary Table S3:** Summary of Ramachandran plot statistics for modelled coat protein structures from four major banana viruses.

| Virus | Total Residues | Gly | Pro | Analyzed | Most Favored (%) | Allowed (%) | Generously Allowed (%) | Disallowed (%) |
| --- | --- | --- | --- | --- | --- | --- | --- | --- |
| BBrMV | 136 | 10 | 5 | 120 | 94.2 | 5.8 | 0.0 | 0.0 |
| BMoV | 217 | 13 | 14 | 188 | 86.2 | 13.8 | 0.0 | 0.0 |
| BSV | 326 | 17 | 9 | 298 | 90.6 | 8.7 | 0.7 | 0.0 |
| BBTV | 175 | 9 | 8 | 156 | 88.5 | 10.9 | 0.6 | 0.0 |

**Supplementary Table S4:** Dock score of interactions between phytochemicals and antiviral agents against coat protein of banana viruses

| Compound | Dock score for binding affinity (kcal/mol) of CP of banana virus | | | |
| --- | --- | --- | --- | --- |
|  | **BBTV** | **BSV** | **BCMV** | **BBrMV** |
| Trigonelline | -4.0 | -4.8 | -4.0 | -3.6 |
| Beta ionone | -5.5 | -5.6 | -5.2 | -5 |
| Cucumerin-A | -8.8 | -7.5 | -6.5 | -9.5 |
| Cucumerin-B | -7.8 | -7.8 | -6.3 | -6.8 |
| 4-Hydroxycinnamic acid | -4.6 | -5.6 | -5.0 | -4.4 |
| Carageenan | -7.9 | -7.7 | -5.9 | -5.3 |
| Acyclovir | -5.6 | -5.5 | -4.4 | -6.1 |
| 5-Azacytidine | -4.9 | -6.5 | -4.9 | -5.0 |
| Cytarabine | -5.9 | -6.4 | -4.8 | -4.5 |
| Ribavirin | -6.6 | -6.6 | -5.2 | -5.3 |
| Ningnanmycin | -8.0 | -6.1 | -8.5 | -5.7 |
| Vidarabine | -6.1 | -6.4 | -5.6 | -4.8 |
| Acycloguanosine | -5.3 | -5.2 | -4.1 | -4.0 |
| 2-Thiouracil | -3.9 | -4.1 | -3.7 | -3.7 |
| Moroxydine hydrochloride | -5.2 | -5.2 | -4.4 | -4.8 |
| Luotonin A | -8.3 | -8.2 | -6.5 | -7.1 |
| Pergularinine | -6.3 | -7.7 | -5.9 | -5.6 |
| Antofine | -6.6 | -7.6 | -6.2 | -5.9 |
| Deoxytylophorinine | -6.9 | -7.8 | -5.9 | -6.1 |
| Pyrroloisoquinoline | -7.4 | -5.8 | -4.3 | -5.8 |
| Streptindole | -7.3 | -8.1 | -5.9 | -5.7 |
| Tryptanthrin | -6.6 | -7.5 | -6.3 | -6.0 |
| Essramycin | -7.3 | -6.8 | -5.5 | -6.4 |
| Chlorogenic acid | -6.9 | -7.1 | -5.7 | -5.7 |
| Peonidin | -7.4 | -6.8 | -6.0 | -6.9 |
| Swertianolin | -8.8 | -9.0 | -5.3 | -6.8 |
| Zidovudine | -6.9 | -8.3 | -5.3 | -6.9 |
| Lutein | -5.5 | -8.0 | -6.6 | -7.2 |
| Isoscoparin | -7.3 | -8.2 | -7.8 | -6.9 |
| Saponin | -9.23 | -11.7 | -8.8 | -13.33 |
| Vicenin-2 | -7.6 | -8.5 | -6.5 | -7 |
| Apigenin -7-o-glucoside | -7.1 | -7.8 | -7.1 | -6.5 |
| Quercetin -3-o-glucoside | -6.9 | -7.1 | -6.2 | -6.9 |
| Isorhamnetin -3-o-glucoside | -7.3 | -7 | -6.1 | -6.4 |
| Kaemferol -3-o-rhamnoside | -7.1 | -7.9 | -6.3 | -6.4 |
| Kaempferol | 5.1 | -7 | -6.1 | -6.6 |
| Myricetin | -7.0 | -7.6 | -5.7 | -6.2 |
| Quercetin | -7.0 | -7.6 | -6.2 | -6.3 |
| (-)-Epigallocatechin gallate | -7.6 | -8 | -6.1 | -6.4 |
| Apigenin | -8.0 | -7.9 | -5.9 | -6.9 |
| Luteolin | -7.5 | -7.3 | -6.3 | -7.1 |
| (-)-Epicatechin | -6.4 | -6.9 | -5.9 | -6 |
| (+)-Gallocatechin | -5.8 | -8.3 | -6.1 | -5.8 |
| Gallocatechin gallate | -7.8 | -8.1 | -6.5 | -5.8 |
| (-)-Epicatechin gallate | -8.0 | -8.0 | -6.1 | -6.9 |
| Epigallocatechin | -6.0 | -6.5 | -5.6 | -6 |
| Procyanidin C1 | -7.6 | -8.4 | -7.5 | -7.5 |
| Procyanidin B4 | -6.9 | -8.5 | -6.8 | -6 |
| Procyanidin B1 | -8.0 | -7.4 | -6.5 | -6.1 |
| Procyanidin B8 | -7.8 | -8.1 | -7.1 | -6.7 |
| Procyanidin B3 | -6.5 | -7.6 | -7.3 | -8.3 |
| Procyanidin B2 | -7.1 | -7.2 | -6.8 | -7.2 |
| Prodelphinidin B | -7.8 | -8.9 | -7.8 | -6.6 |
| Vitexin | -7.5 | -7.6 | -6.1 | -7.1 |
| Isovitexin | -6.9 | -8.3 | -7.8 | -6.3 |
| Orientin | -7.3 | -7.8 | -6.2 | -6.3 |
| Isoorientin | -7.4 | -7.4 | -7.8 | -6.8 |
| Pinoresinol | -6.5 | -7.6 | -6.1 | -6.1 |
| Matairesinol | -5.6 | -6.1 | -5.2 | -6.1 |
| Lariciresinol | -6.1 | -6.9 | -5.9 | -5.2 |
| Secoisolariciresinol | -6.3 | -5.9 | -5.3 | -6.4 |
| 4'-O-Methylirenolone | -6.1 | -6.5 | -6.0 | -6.7 |
| Cucurbitoside K- | -6.2 | -7.1 | -5.2 | -5.8 |
| Cucurbitoside J | -6.7 | -7.0 | -5.5 | -5.5 |
| Cucurbitoside M | -6.4 | -7.9 | -6.3 | -6.5 |
| Cucurbitoside F | -6.4 | -6.9 | -4.7 | -6.1 |
| Cucurbitoside H | -6.4 | -7.4 | -5.2 | -5.5 |
| Cucurbitoside I | -7.4 | -7.6 | -6.1 | -7.5 |
| Cianidanol | -5.4 | -7.3 | -6.3 | -6.6 |
| Gallic acid | -5.9 | -5.5 | -4.5 | -4.3 |
| Bryononic acid | -7.4 | -8.6 | -7.4 | -7.3 |
| p-Coumaraldehyde | -4.6 | -5.3 | -4.3 | -4.7 |
| Irenolone | -6.7 | -7.8 | -5.6 | -6.6 |
| Hydroxyanigorufone | -6.8 | -6.9 | -6.2 | -6.5 |
| Anigorufone | -8.3 | -8 | -5.8 | -7.6 |
| Musanolone F | -7.0 | -7.7 | -7.2 | -6.8 |
| Musanolone E | -6.8 | -7.4 | -6.2 | -6.6 |
| (-)-Musellarin A | -7.2 | -7.2 | -6.3 | -7.1 |
| Musellarin A | -7.7 | -6.2 | -6.1 | -7.2 |

**Supplementary Table S5:** ADMET and drug-likeness profiles of selected antiviral compounds.

| **Properties** | | **441381** | **44257649** | **44588235** | **5858086** |
| --- | --- | --- | --- | --- | --- |
| **A** | Molecular Weight | 594.522 | 552.532 | 443.417 | 436.369 |
|  | Log P | -2.4352 | 1.9491 | -5.8887 | -0.4553 |
|  | Rotatable Bonds | 6 | 5 | 8 | 4 |
|  | Acceptors | 15 | 11 | 12 | 11 |
|  | Doners | 10 | 8 | 8 | 6 |
|  | Water solubility (log mol/L) | -2.724 | -2.914 | -2.175 | -2.623 |
|  | Surface Area | 235.787 | 226.575 | 174.926 | 173.416 |
|  | Caco-2 permeability (in nm/sec) | -0.959 | 0.027 | 0.787 | 0.07 |
|  | Human intestinal absorption (in %) | 8.034 | 54.043 | 8.593 | 49.536 |
|  | Skin Permeability (log K_p_) | -2.735 | -2.735 | -2.735 | -2.735 |
|  | P-glycoprotein substrate | Yes | Yes | No | Yes |
|  | P-glycoprotein I inhibitor | No | Yes | No | No |
|  | P-glycoprotein II inhibitor | No | Yes | No | No |
| **D** | Volume of distribution at steady state (in log L/kg) | 0.564 | -0.138 | 0.156 | 0.234 |
|  | Fraction of unbound plasms | 0.278 | 0.045 | 0.747 | 0.15 |
|  | BBB Permeability (in logBB) | -1.897 | -1.681 | -0.825 | -1.602 |
|  | CNS permeability (in logPS) | -4.784 | -3.967 | -3.994 | -4.087 |
| **M** | CYP2D6 substrate | No | No | No | No |
|  | CYP3A4 substrate | No | Yes | No | No |
|  | CYPIA2 inhibitor | No | No | No | No |
|  | CYP2C19 inhibitor | No | No | No | No |
|  | CYP2C9 inhibitor | No | No | No | No |
|  | CYP2D6 inhibitor | No | No | No | No |
|  | CYP3A4 inhibitor | No | No | No | No |
| **E** | Total Clearance (in log ml/min/kg) | -0.047 | -0.317 | 0.519 | 0.448 |
|  | Renal OCT2 substrate | No | No | No | No |
| **T** | AMES toxicity | No | No | No | No |
|  | Max. tolerated dose (human) | 0.41 | 0.491 | 1.309 | 0.406 |
|  | hERG I inhibitor | No | No | No | No |
|  | hERG II inhibitor | Yes | Yes | No | No |
|  | Oral Rat Acute Toxicity (LD50) | 2.486 | 2.629 | 2.171 | 2.268 |
|  | Oral Rat Chronic Toxicity (LOAEL) | 5.514 | 4.668 | 2.478 | 4.404 |
|  | Hepatotoxicity | No | No | Yes | No |
|  | Skin Sensitisation | No | No | No | No |
|  | *T Pyriformis* toxicity (µg/L) | 0.285 | 0.285 | 0.285 | 0.285 |
|  | Minnow toxicity (log mM) | 11.323 | 5.77 | 6.972 | 4.525 |

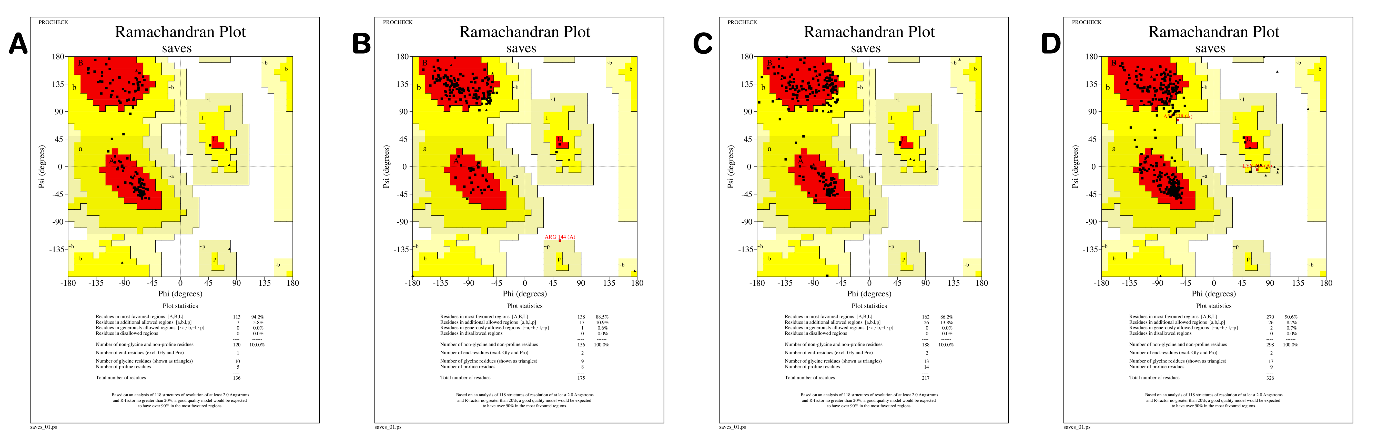

**Supplementary Figure S2:** Ramachandran plot analysis of modelled coat protein structures from major banana viruses.

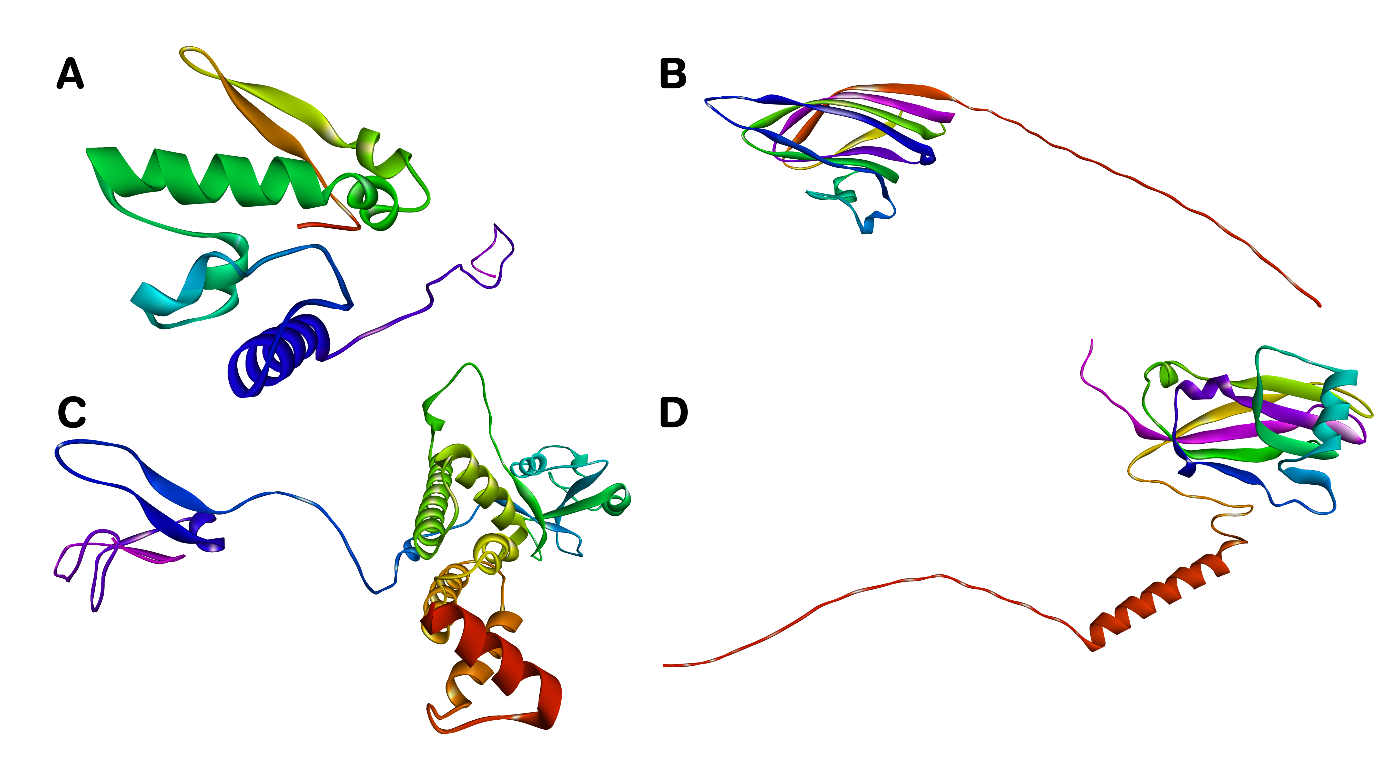

**Supplementary Figure S1:** Predicted three-dimensional structures of coat proteins from four major banana viruses.
